# Can cancer GWAS variants modulate immune cells in the tumor microenvironment?

**DOI:** 10.1101/493171

**Authors:** Yi Zhang, Mohith Manjunath, Jialu Yan, Brittany A. Baur, Shilu Zhang, Sushmita Roy, Jun S. Song

## Abstract

Genome-wide association studies (GWAS) have hitherto identified several genetic variants associated with cancer susceptibility, but the molecular functions of these risk modulators remain largely uncharacterized. Recent studies have begun to uncover the regulatory potential of non-coding GWAS SNPs by using epigenetic information in corresponding cancer cell types and matched normal tissues. However, this approach does not explore the potential effect of risk germline variants on other important cell types that constitute the microenvironment of tumor or its precursor. This paper presents evidence that the breast cancer-associated variant rs3903072 may regulate the expression of *CTSW* in tumor infiltrating lymphocytes. *CTSW* is a candidate tumor-suppressor gene, with expression highly specific to immune cells and also positively correlated with breast cancer patient survival. Integrative analyses suggest a putative causative variant in a GWAS-linked enhancer in lymphocytes that loops to the 3’ end of *CTSW* through three-dimensional chromatin interaction. Our work thus poses the possibility that a cancer-associated genetic variant might regulate a gene not only in the cell of cancer origin, but also in immune cells in the microenvironment, thereby modulating the immune surveillance by T lymphocytes and natural killer cells and affecting the clearing of early cancer initiating cells.

## Main Text

Large-scale genome-wide association studies (GWAS) have been effective in identifying common genetic risk factors for several diseases including cancer. The cancer-associated genetic variants discovered by GWAS, however, are not necessarily causative themselves, but may be in linkage disequilibrium (LD) with other functional variants. Since most GWAS variants are located in non-coding regions, previous functional characterization studies have focused on the gene regulatory function of these linked variants in cancer cells themselves and in matched normal counterparts^1–4^. Although new insights have resulted from these investigations, a provocative question that has not yet been examined is whether select cancer-associated germline variants could also be functional in cell types other than the cell of cancer origin, such as endothelial cells and immune cells, within the heterogeneous tumor microenvironment^5^. For example, in tumor-infiltrating lymphocytes (TIL), genetic variants regulating cytotoxicity-controlling genes may impact TIL’s ability to eliminate cancerous cells, thereby functioning as cryptic modulators of cancer susceptibility that have escaped our attention to date. We have previously introduced a systematic computational framework for studying regulatory functions of noncoding GWAS variants associated with estrogen receptor-positive (ER+) breast cancer^4^; we here apply this approach to present evidence for the possibility that a breast cancer GWAS variant may influence immune cells in the microenvironment of tumor or its precursor.

We hypothesized that the proximity of a genetic variant associated with cancer to those associated with immuno-inflammatory traits might indicate pleiotropy of nearby genes or regulatory variants. In this regard, we examined the NHGRI-EBI GWAS Catalog^6^ and identified rs3903072 as the top breast cancer-associated variant having the highest density of proximal variants within 100kb associated with immuno-inflammatory traits (**Supplementary Methods**; **Supplementary Figure 1**; **Supplementary Table 1**). The single nucleotide polymorphism (SNP) rs3903072 has been found to be associated with ER+ breast cancer in multiple GWAS studies^6–8^, and lies in close physical distance, but with weak genetic linkage (**Supplementary Table 2**), to multiple variants associated with immuno-inflammatory diseases – such as rs118086960 with Psoriasis (an autoimmune disease)^9^, rs77779142 with Rosacea symptom (an inflammatory skin condition)^10^, rs2231884 with inflammatory bowel disease^11^, and rs568617 with Psoriasis and Crohn’s disease (an inflammatory bowel disease)^12; 13^ (**Figure 1A**; **Supplementary Table 1**). A direct link between this non-coding SNP rs3903072 and its regulatory function in mammary epithelial cells is currently unknown; similarly, it remains uncharacterized how and why the aforementioned SNPs in the region affect diverse immuno-inflammatory traits. Discovering the target genes of rs3903072 thus represents a major step towards identifying a potential regulatory mechanism common to both breast cancer susceptibility and immuno-inflammatory traits.

**Figure 1.**
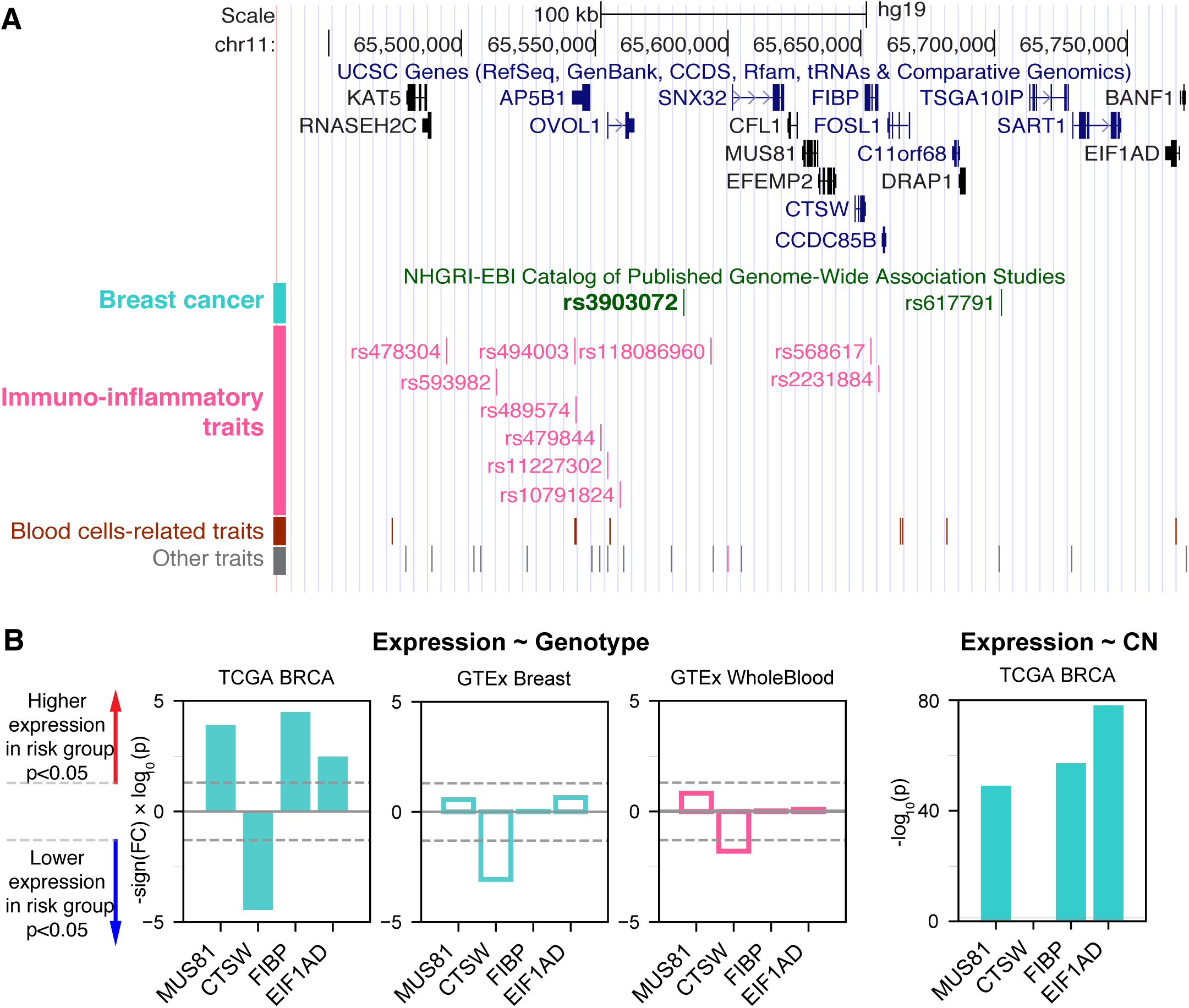
The breast cancer risk variant rs3903072 in the 11q13.1 region. (**A**) The top track shows UCSC genes, and the lower track shows the GWAS variants grouped by their reported traits into four categories (**Supplementary Table 1**). The SNP rs3903072 shows a stronger association with breast cancer (*p* = 2×10^−12^) than the variant rs617791 (*p* = 7×10^−6^). (**B**) The eQTL results for rs3903072. A full list of eQTL genes is in **Supplementary Table 3**. Left three bar plots show the significance of the contribution from rs3903072 genotype to the expression level of *MUS81, CTSW, FIBP* and *EIF1AD*. Positive values represent higher expression as the number of risk allele increases, and negative values represent the opposite trend. *CTSW* is the only eQTL gene with a significant suppression in the risk genotype group. This negative correlation between *CTSW* expression level and the genotype status is confirmed in the three independent datasets shown. The right bar plot shows the significance of the contribution from gene copy number to the expression level of each gene. Filled bar plots represent tumor, while transparent plots represent normal tissues; in this paper, we use the cyan color to indicate breast-related cancer or normal cells, and the magenta color to indicate blood-related cells. The dotted lines in gray mark the significance threshold of *p* = 0.05.

To identify candidate target genes, we applied the approach of expression quantitative trait loci (eQTL)^14^, quantifying the correlation of messenger RNA (mRNA) levels of nearby genes with the genotype status at rs3903072. Using ER+ breast tumor RNA sequencing (RNA-seq) and genotyping data from the Breast Invasive Carcinoma (BRCA) dataset of The Cancer Genome Atlas (TCGA), we identified several significant eQTL genes in *cis* for rs3903072, including *CTSW*, *FIBP*, *MUS81*, and *EIF1AD* (genotype *p*-values of a linear model adjusting for gene copy number: *p* = 3.52×10^−5^, *p* = 3.22×10^−5^, *p* = 1.24×10^−4^, *p* = 3.28×10^−3^, respectively; **Figure 1B**; a complete list of eQTL genes in **Supplementary Table 3**), confirming the results previously reported^7^. Notably, *CTSW* was among the most significant eQTL genes; the negative correlation between *CTSW* expression and the number of risk alleles indicated a tumor-suppressive role of this gene (**Figure 2A**). In line with the eQTL result, survival analysis of BRCA patients showed that higher *CTSW* expression was associated with significantly better survival probability (log-rank *p*-value with median expression cutoff; *p* = 0.026 for ER+ breast cancer patients analyzed in this study (**Figure 2B**); *p* = 7.3×10^−4^ for all BRCA patients in the analysis performed by the Human Protein Atlas (THPA)^15^, image: https://www.proteinatlas.org/ENSG00000172543-CTSW/pathology/tissue/breast+cancer). By contrast, according to the TCGA analysis presented in THPA, other eQTL genes were not significantly associated with breast cancer patient survival.

**Figure 2.**
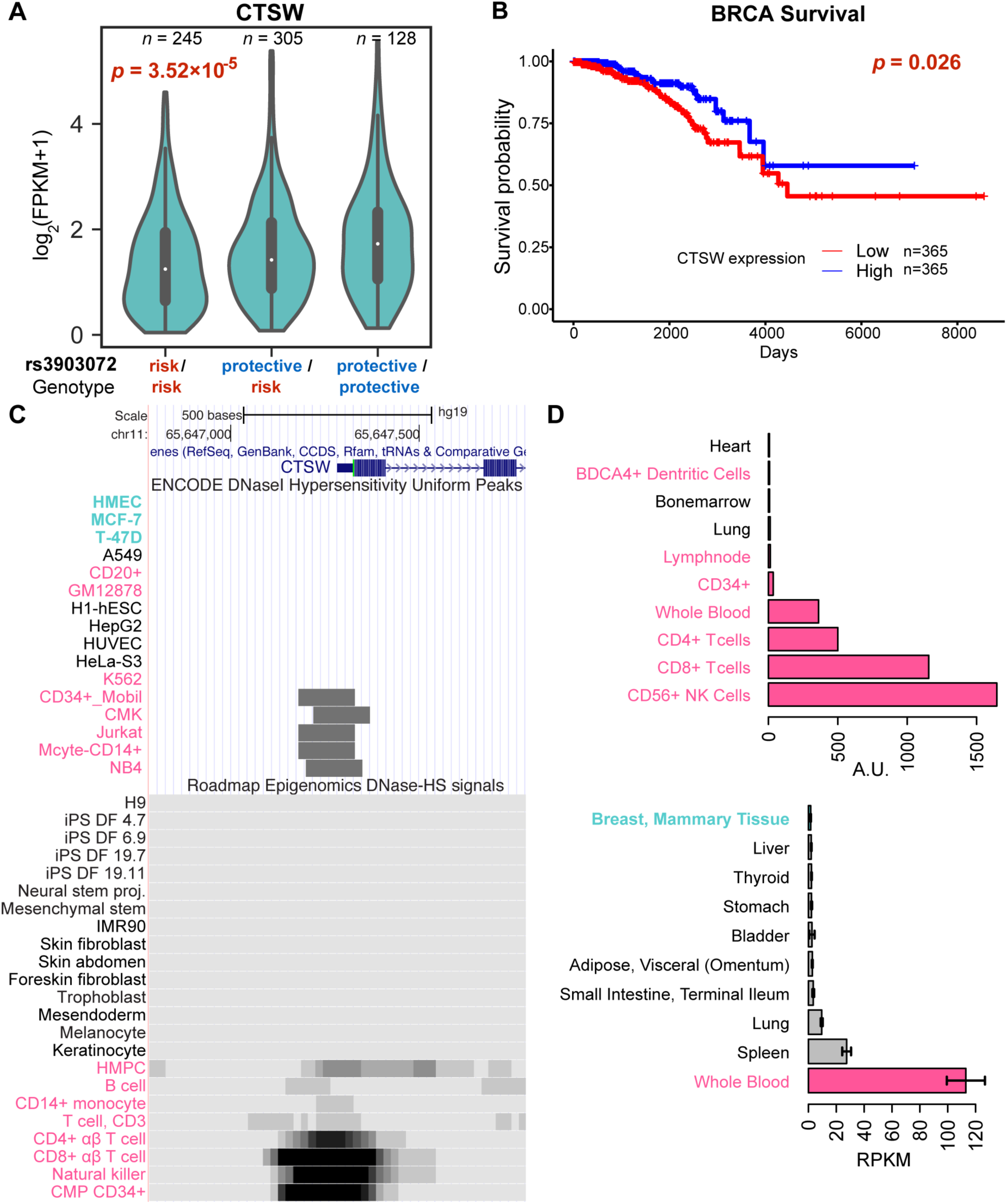
Genotype-dependent suppression and tissue specificity of *CTSW* expression. (**A**) The violin plot of *CTSW* expression levels in three genotype groups at rs3903072, using the TCGA ER+ breast cancer patient data. The *p-*value is for the coefficient of genotype in multivariate linear regression with adjustment for gene copy number. (**B**) Survival analysis of breast cancer patients based on *CTSW* expression levels. The *p-*value is from the log-rank test using the two groups separated by the median expression of *CTSW*. (**C**) The *CTSW* promoter chromatin accessibility in multiple cell lines from various tissue origins. The top ENCODE plot shows the DNase I hypersensitivity (DHS) uniform peaks in ENCODE Tier 1 cell lines, along with the Tier 2/3 cells in which *CTSW* promoter is open. The bottom plot shows the DHS signals for primary cells from Roadmap Epigenomics data. Cells with breast tissue origin are marked as cyan, and blood-related normal or cancer cells are marked as magenta. HMPC, hematopoietic multipotent progenitor cells; CMP CD34+, common myeloid progenitor cells CD34+. (**D**) Distribution of *CTSW* expression in cell lines and in tissues. The top figure shows data from BioGPS, displaying only ten cell types with highest *CTSW* expression. The bottom figure shows data from GTEx, displaying only nine tissue types with highest *CTSW* expression and mammary tissue (ranked 13th).

Consistent with our finding, a similar drop in survival probability with lower *CTSW* expression was also observed in other cancer types including endometrial cancer (Uterine Corpus Endometrial Carcinoma, UCEC) and head and neck cancer (Head-Neck Squamous Cell Carcinoma, HNSC), according to THPA^15^ (log-rank *p*-values with median expression cutoff; UCEC: *p* = 4.1×10^−4^, image: https://www.proteinatlas.org/ENSG00000172543-CTSW/pathology/tissue/endometrial+cancer; HNSC: *p* = 1.9×10^−2^, image: https://www.proteinatlas.org/ENSG00000172543-CTSW/pathology/tissue/head+and+neck+cancer; **Supplementary Methods**). In THPA, although the renal cancer group showed an opposite survival trend when Kidney Renal Clear Cell Carcinoma (KIRC), Kidney Renal Papillary Cell Carcinoma (KIRP) and Kidney Chromophobe (KICH) were combined (THPA *p* = 1.4×10^−4^; log-rank *p*-value with median expression cutoff), the trend was significant only in KIRP when each group was checked separately (THPA; **Supplementary Methods**). Further eQTL analysis confirmed a similar negative correlation between *CTSW* expression level and the rs3903072 risk genotype in UCEC and HNSC, as well as in Low Grade Glioma (LGG), a cancer type not shown in the THPA survival analysis webpage (linear model between expression and rs3903072 genotype; UCEC: *p* = 1.52×10^−3^; HNSC: *p* = 5.45×10^−3^; LGG: *p* = 7.09×10^−3^; **Supplementary Figure 2A**). Together, these results demonstrate that *CTSW* likely has an important biological function in cancer and that the breast cancer risk allele rs3903072-G is significantly associated with decreased expression of *CTSW*.

Interestingly, the following pieces of evidence show that unlike other eQTL genes, *CTSW* is specifically expressed and functions in blood cells, particularly in natural killer (NK) cells and T cells. First, *CTSW* encodes the protein cathepsin W, also named lymphopain, which is a cysteine protease reported to be involved in the cytolytic activity of NK cells and cytotoxic T cells^16; 17^. Second, the *CTSW* promoter region is not accessible in normal mammary cells or breast cancer cells (HMEC, MCF-7, T-47D), but is open in CD8+ T cells, CD56+ NK cells, CD34+ common myeloid progenitor cells, and the acute T cell leukemia cell line Jurkat (**Figure 2C**), according to the DNase I hypersensitive sites sequencing (DNase-seq) data from the Encyclopedia of DNA Elements (ENCODE)^18^ and the Roadmap Epigenomics Project^19^. Third, *CTSW* is predominantly expressed in blood cell lines, but not detectable in human mammary cell lines, according to the gene expression measurements in BioGPS^20^ (microarray; **Figure 2D**) and the Cancer Cell Line Encyclopedia (CCLE^21^, RNA-seq; **Supplementary Figure 3**). Fourth, the *CTSW* promoter is actively transcribed in several lymphocytes, but not in breast cells, according to the cap analysis of gene expression (CAGE) sequencing data from the Functional Annotation of the Mammalian Genome 5 (FANTOM5) project^22^ (**Supplementary Figure 4**). Fifth, among different normal tissue types, *CTSW* expression level measured by Genotype-Tissue Expression (GTEx)^23; 24^ is highest in whole blood, moderate in lung and spleen, and low or undetectable in other tissues, whereas other eQTL genes such as *FIBP*, *MUS81*, and *EIF1AD* are relatively ubiquitously expressed across different tissue types including the mammary gland (**Supplementary Figure 5**). Finally, it has been recently shown that an elevated level of *CTSW* expression is observed in CD8+ T cells with enhanced immunity against bacterial infection and cancer^25^, as well as in renal cancer with high lymphocyte infiltration^26^. Together, these findings demonstrate the high specificity of *CTSW* expression to immune cells, indicating that the *CTSW* mRNA in the TCGA breast tumor bulk RNA-seq has likely arisen from TILs in the heterogeneous tumor microenvironment^5^. In fact, the expression patterns of immune signature genes in TCGA RNA-seq data have been used to infer the abundance of different immune cells in tumor and quantify immune infiltration levels^27^.

**Figure 3.**
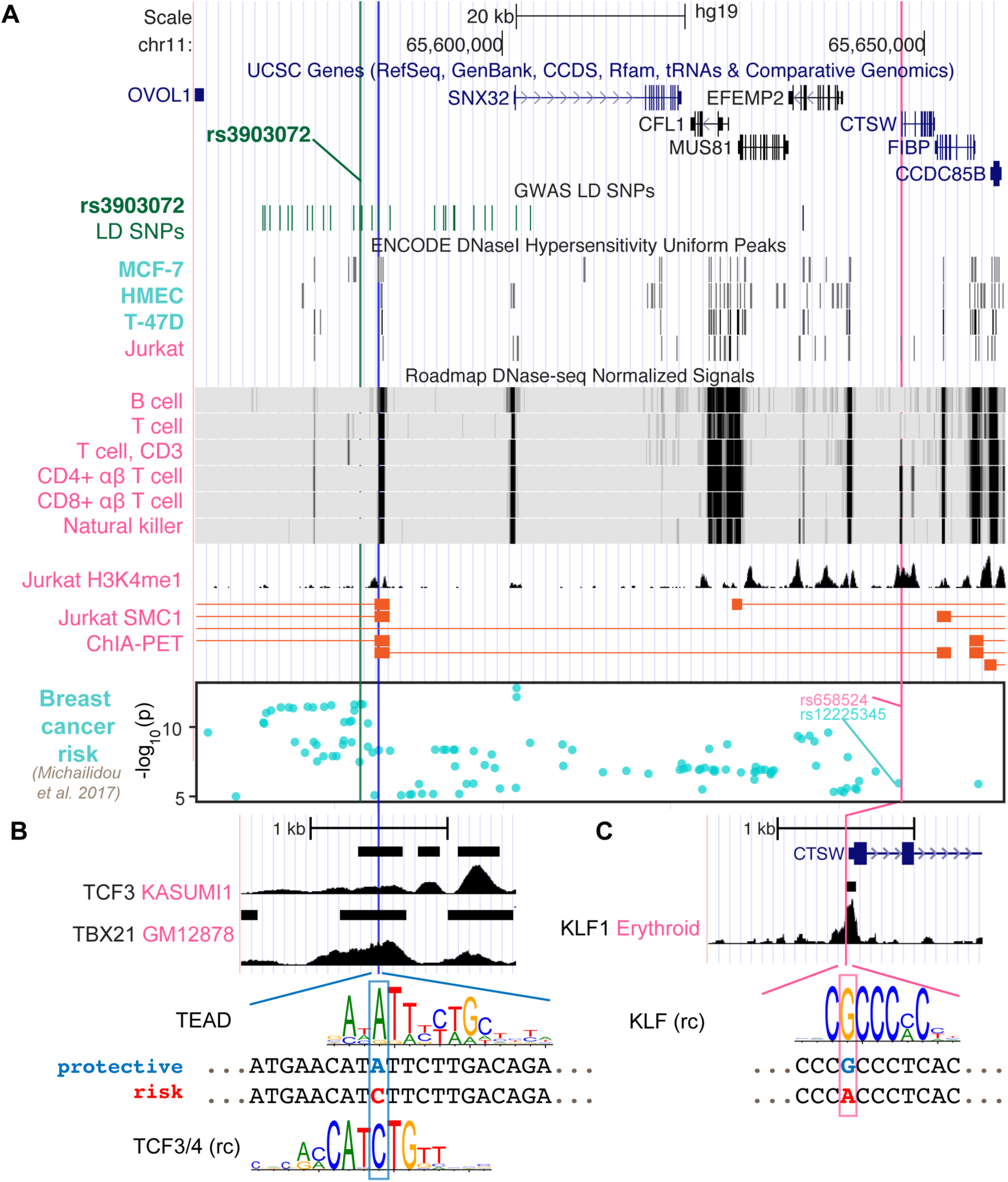
Potential regulatory mechanisms for *CTSW*. (**A**) The genomic region ranging from the GWAS SNP rs3903072 to *CTSW* is shown, including the gene track, the GWAS LD SNPs, epigenetics information and 3D chromatin interaction. The chromatin accessibility data are shown for breast-related cells and blood cells from ENCODE and Roadmap Epigenomics Project. The H3K4me1 modification track and the SMC1 ChIA-PET significant interactions in the Jurkat cell line are from GSE119439 and GSE68978. The bottom Manhattan plot shows the SNPs associated with breast cancer with *p-*value smaller than 10^−5^. Three SNPs are marked: the GWAS SNP rs3903072 and the GWAS-linked putative enhancer SNP in blue, and the *CTSW* promoter SNP in magenta. (**B-C**) Zoomed-in views of the two potential regulatory SNPs. For the GWAS-linked putative enhancer SNP, TCF3 ChIP-seq data in Kasumi1 and TBX21 ChIP-seq data in GM12878 are shown. Two candidate TF motifs predicted to be affected by the SNP are also shown, where the protective allele is marked blue and the risk allele red. For the *CTSW* promoter SNP, KLF1 ChIP-seq profile in Erythroid cells is plotted, along with a KLF family motif (rc: reverse complement).

**Figure 4.**
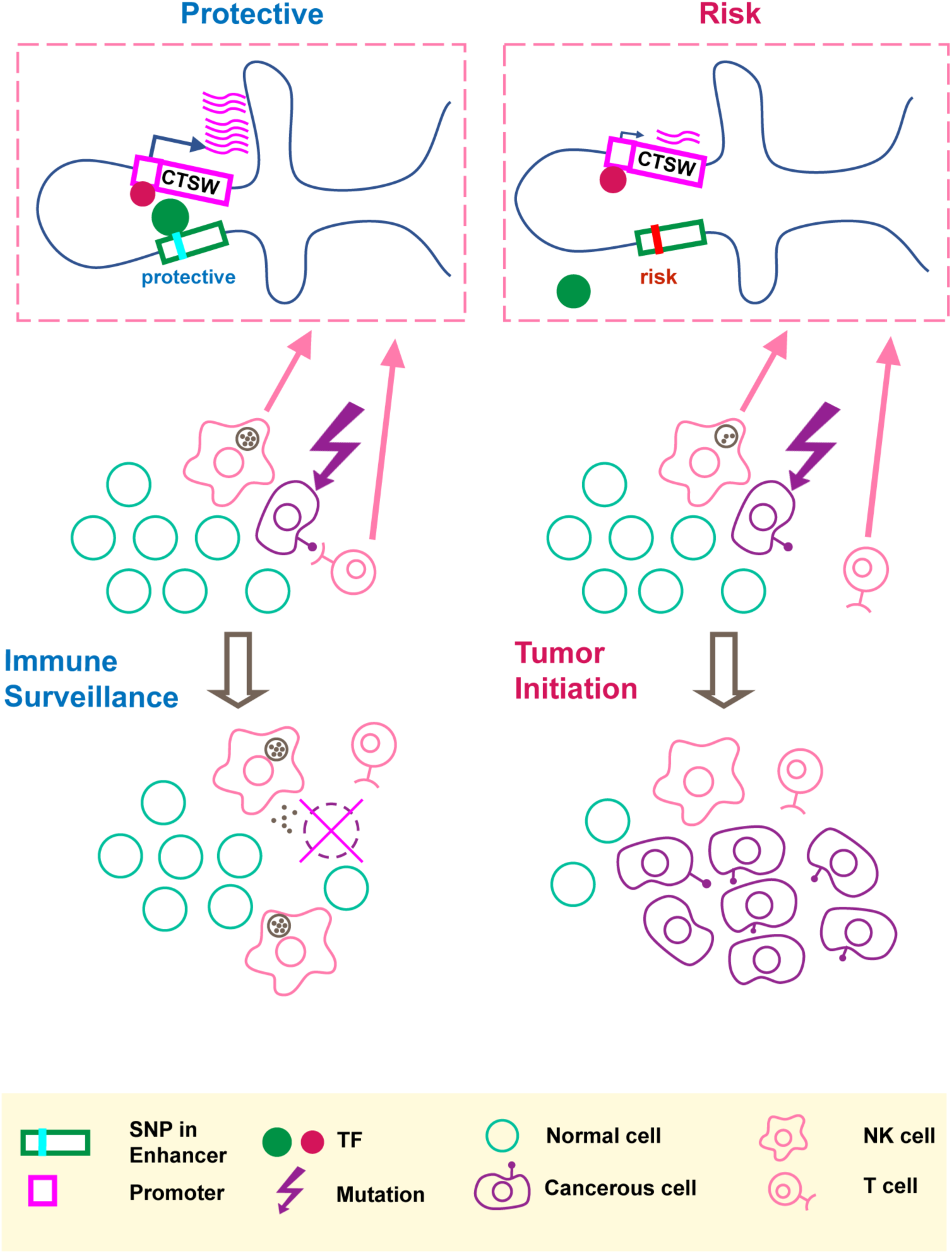
An illustration of our hypothesis. The putative molecular gene-regulation process is shown in the top boxed panels, and the tumor initiation process is illustrated in the two rows below. The left panel shows the process of cancer immune surveillance in people carrying the protective alleles at the GWAS SNP and the predicted functional enhancer SNP. In the top left box, the chromosome carrying the protective alleles produces an abundant *CTSW* mRNA level; as shown here, *CTSW* transcription could be elevated through a transcription activator binding the PRE1 SNP. Note that another scenario, not shown in the illustration, is also possible, where the protective allele could disrupt the binding motif of a transcription repressor. Going back to the cell view, the high level of *CTSW* expression in NK cells or T cells may enhance their cytotoxicity and facilitate their ability to detect and eliminate abnormal cells, such as cancerous mammary epithelial cells that just acquired some oncogenic mutations. This high efficiency of immune surveillance would thus reduce the risk of developing breast cancer. By contrast, the right panel shows the NK/T cells with suppressed *CTSW* expression associated with the risk alleles, resulting in reduced cytotoxic activities and suppressed immune surveillance efficiency, thereby increasing the risk of developing breast cancer.

We thus hypothesized that the breast cancer-associated GWAS variant rs3903072 may regulate *CTSW* in immune cells within the tumor microenvironment, independent of the other eQTL genes that could potentially be regulated separately in breast cancer cells. Several observations supported this idea. First, *CTSW* was the only TCGA-BRCA eQTL gene that remained correlated with the GWAS genotype status in the GTEx normal mammary tissue (*p* = 8.64×10^−4^) and whole blood (*p* = 0.016; **Figure 1B; Supplementary Figure 2B**). Second, *CTSW* was the only eQTL gene that showed no correlation with DNA copy number in TCGA breast cancer data (*p* = 0.72; **Figure 1B**; **Supplementary Table 3**), suggesting regulation unaffected by the genomic amplification or deletion events abundant in cancer cells. Third, *CTSW* was the only eQTL gene showing negative correlation with the number of risk alleles at the GWAS SNP, whereas other eQTL genes had the opposite trend, indicating that *CTSW* may play a tumor-suppressive role in TILs, while others may be involved in promoting cancer progression (linear regression coefficient in BRCA ER+ eQTL: *CTSW*, *r* = −0.22; *FIBP*, *r* = 0.09; *MUS81*, *r* = 0.08; *EIF1AD*, *r* = 0.05; **Figure 1B**). Lastly, *CTSW* was the only gene of known function related to immune cells across the region shown in **Figure 1A** (**Supplementary Table 4**), where multiple GWAS associations point to immuno-inflammatory traits. Even though we do not exclude the possibility that other eQTL genes may also have important functions in TILs or breast cancer cells, the tissue specificity and the correlation structure of *CTSW* expression strongly suggest its significant modulation in tumor-infiltrating immune cells by the GWAS SNP rs3903072 itself or a linked genetic variant.

As the GWAS SNP rs3903072 itself did not reside in an open chromatin region in immune cells (**Figure 3A**), we next searched for putative regulatory variants that could directly control *CTSW* expression. We first found the SNP rs658524 to be located at the center of a DHS peak in *CTSW* promoter among several lymphocyte cell lines (**Supplementary Figure 6A**). On the one hand, the SNP rs658524 was simultaneously linked to two of the immuno-inflammatory GWAS variants. Namely, the GWAS SNP rs77779142, associated with Rosacea symptoms, was in tight LD with the *CTSW* promoter SNP rs658524 (*r*^2^= 0.78 with rs658524; *r*^2^= 0.17 with rs3903072; 1000 Genomes Phase 3 EUR population), despite being closer to the breast cancer GWAS SNP rs3903072 than to rs658524 in genomic distance (16.6 kb to rs3903072 vs. 47.6 kb to rs658524). Another GWAS SNP rs568617, associated with Psoriasis and Crohn’s disease, resided in intron of the gene *FIBP* next to *CTSW* and was in high LD with the *CTSW* promoter SNP rs658524 (*r*^2^ = 0.99 to rs658524; *r*^2^ = 0.19 to rs3903072; **Figure 1A**). On the other hand, the promoter SNP rs658524 was strongly correlated with *CTSW* expression, according to our eQTL analysis in TCGA (linear model between expression and rs658524 genotype; BRCA: *p* = 1.02×10^−17^; UCEC: *p* = 1.50×10^−11^; HNSC: *p* = 1.32×10^−12^; LGG: *p* = 1.43×10^−6^; **Supplementary Figure 7A**) and GTEx (mammary tissue: *p* = 2.19×10^−11^; whole blood: *p* = 8.48×10^−5^; **Supplementary Figure 7B**), consistent with eQTL results from other immune cell studies^28^.

We here note that rs658524-A also showed partial association with breast cancer risk, since the haplotypes carrying the rs658524-A allele were found to be largely biased towards the GWAS risk allele rs3903072-G compared to the alternative allele rs3903072-T, despite the balanced minor allele frequency (MAF) of rs3903072 (rs3903072 MAF=0.46; 188 haplotypes with rs658524-A-rs3903072-G and 3 haplotypes with rs658524-A-rs3903072-T among the 1006 haplotypes from the 1000 Genomes Project Phase 3 EUR population; **Supplementary Figure 6B**). In fact, rs658524 was in weak LD with rs3903072 (low *r*^2^ = 0.186, but high *D* ^′^ = 0.966), with the GWAS risk SNP having a much higher allele frequency than the risk promoter SNP (rs3903072-G frequency: 0.54; rs658524-A frequency: 0.19; 1000 Genomes Project Phase 3, EUR). However, the *CTSW* promoter SNP rs658524 itself did not entirely explain either the GWAS association or the *CTSW* regulation in this region. A recent study reporting GWAS SNPs with a *p-*value less than 10^−5^ for breast cancer^8^ included the *CTSW* locus (Manhattan plot in **Figure 3A**). The top SNP linked to rs658524 was rs12225345 (*r*^2^ = 0.84), which was only moderately associated with breast cancer (*p* = 1.13×10^−6^), separated from the top GWAS signal cluster represented by rs3903072 (*p* = 2.25×10^−12^)^8^. Furthermore, a conditional eQTL analysis showed that within the group of TCGA patients carrying the homozygous genotype rs658524-G/G, the rs3903072 risk allele still displayed a residual negative effect on *CTSW* expression (*Welch* t-test, two-sided, *p* = 6.0×10^−4^; GTEx whole blood data; **Supplementary Figure 8**).

Thus, although the *CTSW* promoter SNP was in high *D* ^′^ with the breast cancer GWAS SNP rs3903072, it did not solely explain the breast cancer risk in 11q13.1, and other functional SNPs likely influenced the expression of *CTSW.* Given the GWAS SNP rs3903072 was located 64 kb away from *CTSW* promoter, we tested whether some putative functional SNPs tightly linked to rs3903072 could affect distal enhancer activities modulating *CTSW* expression. We thus examined all common (*MAF* ≥ 0.05) SNPs from 1000 Genomes Project Phase 3 EUR population in high LD (*r*^2^ ≥ 0.8) with the GWAS SNP rs3903072 and prioritized the potential functional ones by using epigenetic information. In detail, by overlapping the 30 high LD SNPs with DHS of lymphocyte cell lines (**Supplementary Table 5**), we identified three SNP-containing putative regulatory elements (PREs): PRE1 located 3 kb away from rs3903072, PRE2 at *SNX32* promoter, and PRE3 at *EFEMP2* promoter (**Supplementary Figure 9**). Further investigation of the available 3D chromatin interaction data in immune-related cells highlighted PRE1 as the top regulatory element physically interacting with the 3’ end of *CTSW*, as assessed by the Structural Maintenance of Chromosomes protein 1 (SMC1; GSE68978^29^) Chromatin Interaction Analysis by Paired-End Tag Sequencing (ChIA-PET) data (**Figure 3A**). Another Jurkat ChIA-PET data for RAD21, a cohesin complex component, showed an indirect interaction linking PRE1 and *CTSW* mediated through an anchoring element near *EIF1AD*, suggesting multi-way interactions between the several anchoring elements or enhancers (**Supplementary Figure 9**). To contrast the chromatin interaction of PRE1 with *CTSW* between NK/T cells and mammary cells, we predicted high-resolution (5 kb) Hi-C interactions in natural killer cells, CD8+ αβ T cells and benign variant human mammary epithelial cells (vHMEC). Using random forest-based regression models trained separately on high-resolution Hi-C data in five different cell lines^30^ (**Supplementary Methods**), we predicted the contact counts in the three cell types of interest within 1 Mb from rs3903072. Consistent with the ChIA-PET data, natural killer cells were found to have the highest predicted contact count for the pair of rs3903072-PRE1 and *CTSW* in all five models (**Supplementary Figure 10**), in contrast to vHMEC which had the lowest predicted contact counts. These findings together suggested that PRE1 linked to the GWAS SNP could function as a distal regulatory element controlling *CTSW* expression selectively in NK and T cells.

Among the prioritized SNPs residing in the three PREs, we identified rs11227311 in PRE1 as a putative functional SNP (*r*^2^ = 0.89 with rs3903072, 1000 Genomes Phase 3, EUR). More precisely, it not only overlapped a DHS in NK cells, B cells and type 2 T helper cells (ENCODE accession number: ENCFF933OXV; ENCFF772OPR; ENCFF001WTS and ENCFF001WTQ), but also H3K4me1 modification and the ChIA-PET region interacting with *CTSW* 3’ end in Jurkat (**Figure 3A; Supplementary Figure 9**). To identify candidate transcription factors (TF) in PRE1 potentially affected by rs11227311, we scanned the short sequences around the SNP for TF motifs, using the program FIMO (version 4.12.0) and position weight matrices (PWM) collected from multiple motif databases (**Supplementary Methods**). Using our previously described method for measuring the significance of motif disruption by a SNP, based on simulating null mutations on the PWMs^4^, we identified a list of candidate TF motifs disrupted by rs11227311, including the TEAD family, TCF3/4, NR3C1, POU2F1, and ETV5 (**Figure 3B; Supplementary Figure 9**; *p*- values from neutral mutation simulation: *p* = 0.0074, *p* = 0.0086, *p* = 0.002, *p* = 0.013, *p* = 0.045, respectively; **Supplementary Methods**). Although it was difficult to validate which TF can directly bind the PRE1 SNP due to insufficient ChIP-seq data in T/NK cells, we found the PRE1 candidate SNP rs11227311 to be located within a weak TCF3 ChIP-seq peak in Kasumi1 acute myeloid leukemia cell line (GEO GSE43834^31^) (**Figure 3B**). Examination of other ChIP-seq data in ENCODE for TFs in lymphocytes also showed that the SNP rs11227311 is at the center of a strong TBX21 ChIP-seq peak in GM12878 (ENCSR739IHN; **Figure 3B**). TBX21 is a T-box transcription factor controlling important genes in NK cells and type 1 T helper cells^16^, and its binding supports the potential involvement of PRE1 in gene regulation in lymphocytes. Additionally, we performed a motif analysis for the *CTSW* promoter SNP and found that it might disrupt the binding site of the KLF family TFs (*p* = 0.0086; **Supplementary Methods**), the actual binding of which in this region was supported by a KLF1 ChIP-seq dataset in erythroid cells (GSE43625^32^; **Figure 3C**).

In this paper, we performed functional characterization of breast cancer-associated GWAS variants and proposed the idea that a noncoding cancer GWAS SNP might regulate gene expression in immune cells within the tumor microenvironment. **Figure 4** summarizes our hypothesis that the GWAS-linked SNP rs11227311 may directly affect TF binding affinity at the distal enhancer and regulate *CTSW* expression in cytotoxic lymphocytes, thereby affecting their ability to eliminate abnormal cells. As a member in the cathepsin family, *CTSW* is specifically expressed in NK cells and T cells with a potential role in their cytotoxicity; it can also be strongly induced in NK cells by Interleukin-2 (IL-2)^17^ which is a cytokine controlling T cell growth and NK cell cytotoxicity. Although the function of cathepsin W and its precise relation to lymphocyte cytotoxicity remain under debate^33^, the described association between *CTSW* and breast cancer susceptibility renews the interest in this gene as a component of immune surveillance against cancer. Our work highlights the need to examine the effect of GWAS SNPs on gene regulation not only in the cell type of disease, but also in surrounding cells that may modulate the progression of pathology.

## Supporting information

## Acknowledgments

The results appearing here are in part based upon the data generated by the TCGA Research Network (http://cancergenome.nih.gov/, dbGaP accession number phs000424.v6.p1 on 05/06/2016) and the GTEx Project, supported by the Common Fund of the Office of the Director of the National Institutes of Health and by NCI, NHGRI, NHLBI, NIDA, NIMH, and NINDS. We acknowledge the ENCODE consortium that generated the data sets used in the manuscript. Y.Z., M.M., J.Y. and J.S.S. were supported by the NIH 1U54GM114838 grant, awarded by National Institute of General Medical Sciences (NIGMS) through funds provided by the trans-NIH (National Institutes of Health) Big Data to Knowledge (BD2K) initiative, the National Brain Tumor Society and NIH R01CA163336. B.A.B., S.Z, and S.R were supported by the NIH BD2K grant U54 AI117924 and Vilas Fellowship; B.A.B. was also supported by the Genomic Sciences Training Program (NHGRI 5T32HG002760). The content is solely the responsibility of the authors and does not necessarily represent the official views of the National Institutes of Health.

## Declaration of Interests

The authors declare no competing interests.

## Web References

CCLE, https://portals.broadinstitute.org/ccle

dbGaP, http://www.ncbi.nlm.nih.gov/gap

FANTOM5, http://fantom.gsc.riken.jp/5/

GEO, https://www.ncbi.nlm.nih.gov/geo/

GTEx database, https://gtexportal.org/home/

TCGA-GDC, https://portal.gdc.cancer.gov/projects

The Human Protein Atlas, https://www.proteinatlas.org/

