## Supplementary material for "Can cancer GWAS variants modulate immune cells in the tumor microenvironment?"

### SUPPLEMENTARY METHODS

#### *Genome-wide association studies (GWAS) variants*

A list of estrogen receptor-positive (ER+) breast cancer-associated variants were first obtained from *Michailidou et al.*[1] and the NHGRI-EBI GWAS catalog[2]. To identify the GWAS variants associated with immuno-inflammatory traits, we used the NHGRI-EBI GWAS catalog v1.0.2 and the annotation table from the GWAS catalog mapping the GWAS traits to the parent disease category using information from the Experimental Factor Ontology (EFO) database[3]. Using the "Parent term" column, we first selected SNP-trait associations from the GWAS catalog containing the term "Immune system disorder" or "Inflammatory measurement". We also included additional traits in other disease categories with recursive ontology parents containing the keyword "immun" or "inflamm". For example, Crohn' disease was categorized as a digestive system disorder in the GWAS EFO mapping file, but we included it in our study since it is under the "inflammatory bowel disease" category in the EFO ontology database. Finally, we obtained a set of 3404 SNPs associated with immuno-inflammatory traits. For each ER+ breast cancer GWAS SNP, we scanned the proximal region using a moving window of length 100 kb containing the SNP and counted the number of immuno-inflammatory GWAS SNPs within each window. The maximum number of immuno-inflammatory SNPs in these running windows was then recorded for each breast cancer GWAS SNP. When the breast cancer GWAS SNPs were ranked based on these maximum counts, rs3903072 emerged as the top SNP.

A supplementary table from *Michailidou et al.*[4] provides all SNPs associated with breast cancer with $p$-value less than $10^{-5}$. However, as *Michailidou et al.*[4] used the $p$-value threshold of $10^{-8}$ in the final report, we mainly focused on rs3903072 ($p = 2 \times 10^{-12}$) rather than the nearby breast cancer GWAS SNP rs617791 showing a weaker association ($p = 7 \times 10^{-6}$, **Figure 1A**).

***The Cancer Genome Atlas (TCGA) cancer data***

The germline genotypes at tag SNPs of breast cancer (BRCA) patients in the TCGA dataset were downloaded from the TCGA Data Portal. The tumor copy number segmentation data in hg19 from the NCI Genomic Data Commons (GDC) Legacy Archive[5] were used to compute gene copy number (CN). The processed gene expression data in Fragments Per Kilobase Million (FPKM) measured by RNA-seq were downloaded from the TCGA GDC data portal[5]. Germline genotypes from normal tissues and CN/RNA-seq data from tumor tissues were matched using TCGA barcodes representing patients. For three other TCGA cancer datasets – Uterine Corpus Endometrial Carcinoma (UCEC), Head-Neck Squamous Cell Carcinoma (HNSC) and Low

Grade Glioma (LGG) – only the germline genotypes and processed gene expression levels were downloaded.

***The Genotype-Tissue Expression (GTEx) project data***

The GTEx[6; 7] gene expression levels in RPKM (Reads Per Kilobase Million) were downloaded from the GTEx portal (file: GTEx_Analysis_v6_RNA-seq_RNA-SeQCv1.1.8_gene_rpkm.gct), and the annotations for the samples and tissues were obtained from GTEx_Data_V6_Annotations_SampleAttributesDS.txt. We analyzed the GTEx data with two aims: comparing the expression levels of a certain gene across different tissues, and analyzing the correlation between the genotype at a GWAS SNP and candidate target gene expression level. For the first purpose, we used the mean expression level of *CTSW* measured in the GTEx tissues (**Figure 2D**). The tissue-wise distributions of other nearby genes were also shown using the UCSC GTEx track[8] (**Supplementary Figure 5**). For the second aim, we used the fully processed, filtered and normalized gene expression levels in the breast mammary tissue and the whole blood tissue from GTEx Analysis V7 (dbGaP Accession phs000424.v7.p2); the imputed genotypes were extracted from the controlled-access dbGaP Accession phg000520.v2 (GTEx V2) dataset.

***Genotype imputation for TCGA datasets***

For genotype imputation in BRCA, UCEC, HNSC and LGG datasets, raw genotypes with genotype confidence score greater than 0.1 in birdseed format were marked as missing genotypes to be imputed along with the non-probed SNPs. The filtered genotypes were then uploaded to the Michigan Imputation Server[9] for genotype imputation, selecting the Haplotype Reference

Consortium (HRC) r1.1 2016[10] as a reference panel, Eagle v2.3[11] for phasing, and EUR population as the quality control option.

***Expression quantitative trait loci (eQTL) analysis for TCGA-BRCA***

In this paper, a pilot eQTL analysis was first performed among ER+ breast cancer patients. For this ER+ breast cancer analysis, we constructed a multivariate linear model, regressing the gene expression levels against the genotypes at the GWAS SNP rs3903072 as well as the gene CN[12]. The gene expression levels in FPKM were log-transformed as $log_2(FPKM + 1)$ and taken to be the response. The SNP genotypes were encoded as the number of risk alleles (0, 1 or 2), and the gene CN was computed by taking gene length-weighted average of tumor CN segmentation data, transforming the segmentation unit back to the copy number unit ($CN = 2 \times 2^{segmentation}$). Multivariate linear regression was then performed for each gene within the 3 Mb region centered at rs3903072. Genes with $\overline{FPKM} \geq 1$ ($\overline{FPKM}$: mean expression among tumor samples) and the genotype $p$-value $\leq 0.05$ from the linear regression were selected for further investigation. Among them, *CTSW* was identified as a signal different from other genes, as described in the main manuscript; the three significant eQTL genes near *CTSW* – *FIBP, MUS81 and EIF1AD* – were then selected for comparison with *CTSW* in **Figure 1B**, because of their strong eQTL correlation that was also observed in an earlier study[1].

A second stage of eQTL analyses was performed to validate the genotype correlation of *CTSW* in other cancer types. For these eQTL analyses using BRCA, UCEC, HNSC LGG and GTEx data, linear regression models between *CTSW* expression and the genotype status at rs3903072 were constructed. All RNA-seq FPKM values from TCGA were log-transformed as in BRCA. For

GTEx, the normalized gene expression matrices of whole blood and breast mammary tissues were downloaded from GTEx Analysis V7 release (dbGaP Accession phs000424.v7.p2).

#### *TCGA survival analysis*

Survival analysis in TCGA ER+ breast cancer patients was performed using the clinical data obtained from TCGA-GDC. The differences in survival rate between the two breast cancer patient groups separated by *CTSW* median expression level was tested using log-rank test with the R packages survival[13] and survminer[14]. Survival analysis results were also obtained in endometrial cancer (UCEC), head and neck cancer (HNSC), and renal cancer, all from the human protein atlas (THPA)[15], choosing the median expression level as the cutoff threshold for grouping patients (UCEC: https://www.proteinatlas.org/ENSG00000172543-CTSW/pathology/tissue/endometrial+cancer; HNSC: https://www.proteinatlas.org/ENSG00000172543-CTSW/pathology/tissue/head+and+neck+cancer; Renal cancer: https://www.proteinatlas.org/ENSG00000172543-CTSW/pathology/tissue/renal+cancer). The three datasets in renal cancer were checked separately, including Kidney Renal Clear Cell Carcinoma (KIRC; https://www.proteinatlas.org/ENSG00000172543-CTSW/pathology/tissue/renal+cancer/KIRC), Kidney Renal Papillary Cell Carcinoma (KIRP; https://www.proteinatlas.org/ENSG00000172543-CTSW/pathology/tissue/renal+cancer/KIRP), and Kidney Chromophobe (KICH; https://www.proteinatlas.org/ENSG00000172543-CTSW/pathology/tissue/renal+cancer/KICH).

#### *CTSW expression and promoter transcription activity*

*CTSW* gene expression in a variety of tissues and cell lines was obtained from BioGPS GeneAtlas[16], cancer cell line encyclopedia (CCLE)[17] and functional annotation of the mammalian genome (FANTOM)[18] resources. Microarray gene expression data for *CTSW* were directly downloaded from the BioGPS web resource. There were 176 samples including replicates, and the mean expression of replicates was calculated for each tissue. CCLE gene expression data in RPKM units from RNA-seq data of 1156 samples were downloaded from the CCLE website (file: CCLE_DepMap_18q3_RNAseq_RPKM_20180718.gct). FANTOM gene expression data in TPM units was obtained from the FANTOM web resource (file: hg19.gene_phase1and2combined_tpm.osc.txt). The FANTOM data consisted of 1829 samples (tissue and cell type information obtained from HumanSamples2.0.sdrf.xlsx).

***Chromatin accessibility at CTSW promoter***

DNase-seq chromatin accessibility measurements in various tissues were obtained from Encyclopedia of DNA Elements (ENCODE)[19] and the Roadmap Epigenomics project[20]. For the ENCODE data, the DNase I hypersensitivity sites (DHS) tracks of all cell types in ENCODE Tier 1 were displayed, together with the three cell lines related to breast tissue (HMEC, MCF-7, T-47D). We also examined all cell types in ENCODE Tier 2 and Tier 3, selecting the ones with a DHS at *CTSW* promoter to include in **Figure 2C**. For the Roadmap Epigenomics data, we displayed the wiggle track of the first DNase-seq replicate in each cell type (**Figure 2C**).

***ChIA-PET data analysis***

We searched the ENCODE and Gene Expression Omnibus (GEO)[21] databases for available three-dimension (3D) chromatin interaction data in lymphocyte cell lines expressing *CTSW*, and found two ChIA-PET datasets in the Jurkat cell line for the proteins SMC1 (GSE68978[22]) and RAD21 (ENCODE Accession ENCSR361AYD). For the SMC1 ChIA-PET data, we used the significant interactions processed and merged by the authors from GEO. For the RAD21 ChIA-PET data, we collected all the raw sequences and generated the chromatin interactions using ChIA-PET 2[23] with default parameters.

***Random forest regression approach for predicting high-resolution chromatin contact counts***

To predict high-resolution Hi-C interactions around the GWAS SNP rs3903072 in Natural Killer cells, T cells and vHMEC, we trained a local random forest regression model within1 Mb of the SNP, using an approach similar to our previously published method[24]. vHMEC serves as a control cell line, since it is a homogeneous, non-cancerous mammary epithelial cell line and does not contain other cell types such as T cells and NK cells. We trained the models on region-pairs involving the chr11:65580000-65585000 (hg19) 5 kb bin, which overlaps the GWAS SNP and PRE1, and 5 kb bins within 1 Mb from the SNP, using published high-resolution (5 kb) Hi-C datasets in five different cell lines[25] and complementary one-dimensional signals as features; these signals were histone marks, DNase I and DNase I accessible sequence specific motifs for CTCF, RAD21 and TBP[26]. Histone marks and DNase I data were obtained from ENCODE and the Roadmap Epigenomics Project for the five training (GM12878, K562, HUVEC, NHEK, HMEC) and three test (NK, T cells and vHMEC) cell lines. Since histone datasets for all the features were not available in vHMEC, we used imputed signals from the Avocado pipeline[27]. Data processing and normalization were done as described in Zhang *et al.*[24] and included normalization for sequencing depth and collapsing replicates by median. To account for overall differences in signal across cell lines, we additionally discretized each of 5 kb ChIP-seq signals using k-means clustering with k=20.

A region was represented as a 10-dimensional feature vector, each dimension corresponding to one of the 10 genome-wide datasets (6 histone ChIP-seq, DNase-seq and 3 DNase-seq derived motifs). Features for a pair of regions were obtained by concatenating the 10-dimensional feature vectors of the two regions together with the feature vector of the intervening region between the two regions and the distance between the two regions to obtain a feature vector of size 31. The feature associated with the intervening region was the mean signal value of the features in the region for ChIP-seq and DHS, similar to the "WINDOW" feature in TargetFinder[28]. Once trained, we used the models to generate contact count predictions in the 1 Mb region in NK cells, CD8+ αβ T cells and vHMEC, using feature datasets from the Roadmap Epigenomics database.

***Prioritization of functional SNPs linked to GWAS SNPs***

We first selected all common ($MAF \geq 0.05$) SNPs from 1000 Genomes Project Phase 3 in high linkage disequilibrium (LD) ($r^2 \geq 0.8$) with rs3903072. All LD values were calculated using the haplotypes from the 1000 Genomes Project Phase 3 EUR population, since the original GWAS SNP rs3903072 was discovered in the European population. To prioritize SNPs located in putative regulatory regions (PREs), we collected DHS from ENCODE and the Roadmap Epigenomics project in lymphocytes-related cells, such as T cells, NK cells, B cells, T helper cells and common myeloid progenitor cells (**Supplementary Table 5**). The LD SNPs overlapping any of the DHS peaks were prioritized for further investigation. In addition to DHS, we also used H3K4me1 modification (processed wiggle track in Jurkat cells from GSE119439[29]) to prioritize SNPs within putative regulatory elements.

***Motif analysis***

TF position-specific weight matrices (PWM) were collected from HOCOMOCO Human v10[30], FACTORBOOK[31; 32], TRANSFAC[33], JASPAR vertebrates[34] and Jolma2013[35]. To identify potential binding sites affected by SNPs, we used the program FIMO[36] (version 4.12.0) to scan the 51 bp sequences carrying either allele of each prioritized SNP in the center (FIMO threshold $10^{-3}$). The statistical significance of motif disruption or creation effect of the SNP alleles was then measured using our previous method of simulating null mutations in motif sequences[12]. Motif logos were plotted using WebLogo3[37] based on PWMs or their reverse complement matrices.

***ChIP-seq analysis***

ChIP-seq data for relevant TFs were collected from ENCODE and GEO. Processed wiggle tracks and peaks were downloaded and presented when available in hg19 (Jurkat H3K4me1 ChIP-seq track from GSE119439[29]; TBX21 ChIP-seq pooled wiggle track and peaks in GM12878 from ENCODE ENCFF193RDB and ENCFF869HSY). For TCF3 ChIP-seq data in Kasumi1 and KLF1 ChIP-seq in GM12878, the raw sequences were downloaded from GSE43834[38] and GSE43625[39], respectively, and processed using SRA Toolkit[40], mapped to hg19 using BWA[41] ( -n 2), and analyzed for peaks using MACS2[42] (callpeak: -q 0.1 --SPMR; bdgcmp: -m FC).

### SUPPLEMENTARY TABLES

**Supplementary Table 1.** GWAS traits around rs3903072 for the region shown in **Figure 1A**.

| GWAS SNP ID | Trait Category | GWAS traits reported in each study | PUBMED ID for each study |
|---|---|---|---|
| rs10750766 | Blood cells-related | Diastolic blood pressure x alcohol consumption interaction (2df test) ; High light scatter reticulocyte count ; High light scatter reticulocyte percentage of red cells ; Immature fraction of reticulocytes ; Systolic blood pressure x alcohol consumption interaction (2df test) | 29912962 ; 27863252 ; 27863252 ; 27863252 ; 29912962 |
| rs10791824 | Immuno-inflammatory | Atopic dermatitis | 26482879 |
| rs10896045 | Blood cells-related | Blood protein levels | 29875488 |
| rs11227302 | Immuno-inflammatory | Systemic lupus erythematosus | 28714469 |
| rs11227306 | Other traits | DNA methylation (variation) | 23725790 |
| rs11602769 | Other traits | Allergic sensitization | 30013184 |
| rs11604462 | Other traits | Glomerular filtration rate (creatinine) | 28452372 |
| rs118086960 | Immuno-inflammatory | Psoriasis | 28537254 |
| rs12223803 | Blood cells-related | Albumin-globulin ratio | 29403010 |
| rs12576766 | Other traits | Serum uric acid levels | 29403010 |
| rs185542523 | Other traits | Maximum cranial width | 29698431 |
| rs201316070 | Other traits | Systolic blood pressure x smoking status (current vs non-current) interaction (2df test) ; Systolic blood pressure x smoking status (ever vs never) interaction (2df test) | 29455858 ; 29455858 |
| rs2231884 | Immuno-inflammatory | Inflammatory bowel disease | 23128233 |
| rs3825068 | Blood cells-related | Blood protein levels | 29875488 |
| rs3903072 | Breast cancer | Breast cancer ; Breast cancer ; Breast cancer | 23535729 ; 25751625 ; 29059683 |
| rs4014195 | Other traits | Chronic kidney disease ; Glomerular filtration rate (creatinine) | 20383146 ; 26831199 |
| rs478304 | Immuno-inflammatory | Acne (severe) ; Spherical equivalent or myopia (age of diagnosis) | 24927181 ; 29808027 |

| | | | |
|---|---|---|---|
| rs479844 | Other traits | Allergic disease (asthma, hay fever or eczema) ; Atopic dermatitis ; Atopic dermatitis ; Atopic march | 29083406 ; 22197932 ; 26482879 ; 26542096 |
| rs489574 | Immuno-inflammatory | Systemic lupus erythematosus | 28714469 |
| rs494003 | Immuno-inflammatory | Systemic lupus erythematosus ; Systemic lupus erythematosus | 26502338 ; 27399966 |
| rs526631 | Blood cells-related | Eosinophil percentage of granulocytes ; Neutrophil percentage of granulocytes | 27863252 ; 27863252 |
| rs568617 | Immuno-inflammatory | Chronic inflammatory diseases (ankylosing spondylitis, Crohn's disease, psoriasis, primary sclerosing cholangitis, ulcerative colitis) (pleiotropy) ; Crohn's disease | 26974007 ; 26192919 |
| rs5792377 | Other traits | Heel bone mineral density | 30048462 |
| rs593982 | Immuno-inflammatory | Atopic dermatitis | 23042114 |
| rs617791 | Other traits | Breast cancer | 29059683 |
| rs634534 | Blood cells-related | Eosinophil counts ; Sum eosinophil basophil counts | 27863252 ; 27863252 |
| rs637571 | Blood cells-related | Eosinophil percentage of white cells | 27863252 |
| rs642803 | Other traits | Educational attainment (MTAG) ; Highest math class taken (MTAG) ; Urate levels | 30038396 ; 30038396 ; 23263486 |
| rs7123489 | Other traits | Creatinine levels | 29403010 |
| rs72941051 | Other traits | Diastolic blood pressure x smoking status (current vs non-current) interaction (2df test) ; Diastolic blood pressure x smoking status (ever vs never) interaction (2df test) ; Systolic blood pressure x smoking status (current vs non-current) interaction (2df test) ; Systolic blood pressure x smoking status (ever vs never) interaction (2df test) | 29455858 ; 29455858 ; 29455858 ; 29455858 |
| rs77291001 | Other traits | Maximum cranial width | 29698431 |
| rs77779142 | Immuno-inflammatory | Rosacea symptom severity* | 29771307 |
| rs9795139 | Other traits | Serum uric acid levels | 29403010 |

* rs77779142 is not shown in **Figure 1A**, because Rosacea symptom was not listed as an immune-related disease in the original GWAS annotation; it is recorded here as an immuno-inflammatory variant, since Rosacea is an inflammatory skin condition.

**Supplementary Table 2**. Genetic linkage between rs3903072 and the nearby immuno-inflammatory variants.

| Immuno-inflammatory SNP | $r^2$ | D' | Immuno-inflammatory risk allele | Immuno-inflammatory risk allele frequency | Allele correlated with rs3903072-G (risk) |
|---|---|---|---|---|---|
| rs478304 | 0.0598 | 0.2474 | T | 0.534 | Linkage Equilibrium |
| rs593982 | 0.0227 | 0.4476 | C | 0.883 | Linkage Equilibrium |
| rs494003 | 0.1078 | 0.737 | A | 0.189 | A |
| rs489574 | 0.0003 | 0.0226 | A | 0.343 | Linkage Equilibrium |
| rs479844 | 0.3262 | 0.6931 | G | 0.557 | A |
| rs11227302 | 0.1057 | 0.7416 | A | 0.184 | A |
| rs10791824 | 0.3234 | 0.7157 | G | 0.575 | A |
| rs118086960 | 0.0233 | 0.1553 | T | 0.532 | Linkage Equilibrium |
| rs77779142 | 0.1673 | 1.0000 | T | 0.164 | T |
| rs568617 | 0.1851 | 0.9657 | T | 0.189 | T |
| rs2231884 | 0.1586 | 0.9737 | T | 0.164 | T |

**Supplementary Table 3.** List of eQTL genes correlated with the rs3903072 genotype within 3 Mb of the SNP.

Available as a separate supplementary file.

**Supplementary Table 4.** Annotation of genes in the rs3903072-*CTSW* region from PANTHER[43].

| Gene ID | Mapped IDs | Gene Name; Gene Symbol | PANTHER Family/Subfamily | PANTHER Protein Class |
|---|---|---|---|---|
| HUMAN\|HGNC=25104\|UniProtKB=Q2VPB7 | AP5B1 | AP-5 complex subunit beta-1;AP5B1 | AP-5 COMPLEX SUBUNIT BETA-1 (PTHR34033:SF1) | |
| HUMAN\|HGNC=24144\|UniProtKB=Q96A11 | GAL3ST3 | Galactose-3-O-sulfotransferase 3;GAL3ST3 | GALACTOSE-3-O-SULFOTRANSFERASE 3 (PTHR14647:SF76) | |
| HUMAN\|HGNC=13718\|UniProtKB=P15407 | FOSL1 | Fos-related antigen 1;FOSL1 | FOS-RELATED ANTIGEN 1 (PTHR23351:SF6) | basic leucine zipper transcription factor(PC00056) |
| HUMAN\|HGNC=5275\|UniProtKB=Q92993 | KAT5 | Histone acetyltransferase KAT5;KAT5 | HISTONE ACETYLTRANSFERASE KAT5 (PTHR10615:SF124) | acetyltransferase(PC00038); chromatin/chromatin-binding protein(PC00077);zinc finger transcription factor(PC00244) |
| HUMAN\|HGNC=26423\|UniProtKB=Q86XE0 | SNX32 | Sorting nexin-32;SNX32 | SORTING NEXIN-32 (PTHR45850:SF3) | |
| HUMAN\|HGNC=3705\|UniProtKB=O43427 | FIBP | Acidic fibroblast growth factor intracellular-binding protein;FIBP | ACIDIC FIBROBLAST GROWTH FACTOR INTRACELLULAR-BINDING PROTEIN (PTHR13223:SF2) | |

| | | | | |
|---|---|---|---|---|
| HUMAN\|HGNC=26555\|UniProtKB=Q3SY00 | TSGA10IP | Testis-specific protein 10-interacting protein;TSGA10IP | TESTIS-SPECIFIC PROTEIN 10-INTERACTING PROTEIN (PTHR21501:SF5) | |
| HUMAN\|HGNC=10769\|UniProtKB=Q13435 | SF3B2 | Splicing factor 3B subunit 2;SF3B2 | SPLICING FACTOR 3B SUBUNIT 2 (PTHR12785:SF6) | |
| HUMAN\|HGNC=2478\|UniProtKB=Q15828 | CST6 | Cystatin-M;CST6 | CYSTATIN-M (PTHR47033:SF1) | |
| HUMAN\|HGNC=2546\|UniProtKB=P56202 | CTSW | Cathepsin W;CTSW | CATHEPSIN W (PTHR12411:SF101) | cysteine protease(PC00081);protease inhibitor(PC00191) |
| HUMAN\|HGNC=10538\|UniProtKB=O43290 | SART1 | U4/U6.U5 tri-snRNP-associated protein 1;SART1 | U4/U6.U5 TRI-SNRNP-ASSOCIATED PROTEIN 1 (PTHR14152:SF5) | extracellular matrix protein(PC00102) |
| HUMAN\|HGNC=8525\|UniProtKB=O14753 | OVOL1 | Putative transcription factor Ovo-like 1;OVOL1 | TRANSCRIPTION FACTOR OVO-LIKE 1-RELATED (PTHR10032:SF217) | |
| HUMAN\|HGNC=17116\|UniProtKB=Q8NEC5 | CATSPER1 | Cation channel sperm-associated protein 1;CATSPER1 | CATION CHANNEL SPERM-ASSOCIATED PROTEIN 1 (PTHR47193:SF1) | |
| HUMAN\|HGNC=24116\|UniProtKB=Q8TDP1 | RNASEH2C | Ribonuclease H2 subunit C;RNASE | RIBONUCLEASE H2 SUBUNIT C (PTHR47063:S | |

| | | | | |
|---|---|---|---|---|
| | | H2C | F1) | |
| HUMAN\|HGNC=1874\|UniProtKB=P23528 | CFL1 | Cofilin-1;CFL1 | COFILIN-1 (PTHR11913:SF17) | non-motor actin binding protein(PC00165) |
| HUMAN\|HGNC=2482\|UniProtKB=P04080 | CST6 | Cystatin-B;CSTB | CYSTATIN-B (PTHR11414:SF22) | cysteine protease inhibitor(PC00082) |
| HUMAN\|HGNC=17397\|UniProtKB=O75531 | BANF1 | Barrier-to-autointegration factor;BANF1 | BARRIER-TO-AUTOINTEGRATION FACTOR (PTHR12912:SF10) | |
| HUMAN\|HGNC=30032\|UniProtKB=Q6VY07 | PACS1 | Phosphofurin acidic cluster sorting protein 1;PACS1 | PHOSPHOFURIN ACIDIC CLUSTER SORTING PROTEIN 1 (PTHR13280:SF16) | |
| HUMAN\|HGNC=28147\|UniProtKB=Q8N9N8 | EIF1AD | Probable RNA-binding protein EIF1AD;EIF1AD | RNA-BINDING PROTEIN EIF1AD-RELATED (PTHR21641:SF0) | |
| HUMAN\|HGNC=3219\|UniProtKB=O95967 | EFEMP2 | EGF-containing fibulin-like extracellular matrix protein 2;EFEMP2 | EGF-CONTAINING FIBULIN-LIKE EXTRACELLULAR MATRIX PROTEIN 2 (PTHR24034:SF96) | annexin(PC00050);calmodulin(PC00061);cell adhesion molecule(PC00069);extracellular matrix glycoprotein(PC00100);extracellular matrix structural protein(PC00103);signaling molecule(PC00207) |
| HUMAN\|HGNC=28801\|UniProtKB=Q9H3H3 | C11orf68 | UPF0696 protein C11orf68; C11orf68 | UPF0696 PROTEIN C11ORF68 (PTHR31977:SF1) | |

| | | | |
|---|---|---|---|
| HUMAN\|HGNC=29814\|UniProtKB=Q96NY9 | MUS81 | Crossover junction endonuclease MUS81;MUS81 | CROSSOVER JUNCTION ENDONUCLEASE MUS81 (PTHR13451:SF0) |

**Supplementary Table 5.** List of DNase-seq peak files for lymphocytes.

| **Cell line (treatment)** | **ENCODE accessions numbers of DHS peak files** |
|---|---|
| Cd4+, helper T cell | ENCFF569GSL, ENCFF907KBL, ENCFF988HSM |
| Cd8+, alpha-beta T cell | ENCFF071RMN, ENCFF422HNI, ENCFF662NTN |
| Jurkat clone E61 | ENCFF582KJR, ENCFF837AOM |
| Natural killer cell | ENCFF224LJW, ENCFF933OXV |
| T cell | ENCFF026SFK, ENCFF286UIJ, ENCFF304TBE, ENCFF345YDG, ENCFF923EVD |
| B cell | ENCFF210RAG, ENCFF654IWG, ENCFF772OPR |
| T helper17 cell | ENCFF001WCL, ENCFF001WTC |
| T helper1 cell | ENCFF434LIX, ENCFF001WCS, ENCFF001WTE, ENCFF001WTI, ENCFF001WTL, ENCFF773IYG, ENCFF001WCQ, ENCFF001WTF, ENCFF001WTM |
| T helper2 cell | ENCFF570MGY, ENCFF001WCW, ENCFF001WTO, ENCFF001WTS, ENCFF001WTU, ENCFF001WCU, ENCFF001WTQ |
| Common myeloid progenitor, CD34+ | ENCFF037XOG, ENCFF387SIU, ENCFF401NSY, ENCFF457SNT, ENCFF479XZN, ENCFF499OEI, ENCFF600EJV, ENCFF664WUJ, ENCFF686ZXP, ENCFF727NEX, ENCFF770BVB, ENCFF918ICP, ENCFF182JTX, ENCFF264UIE, ENCFF927MCZ |

### SUPPLEMENTARY FIGURES

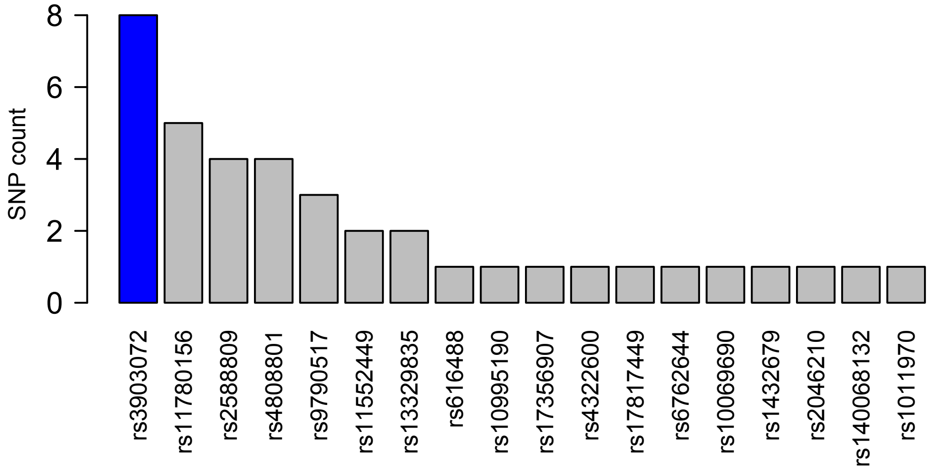

**Supplementary Figure 1.** Number of GWAS variants associated with immuno-inflammatory traits around each ER+ breast cancer GWAS SNP. Only those breast cancer GWAS SNPs with a non-zero count of immuno-inflammatory SNPs within +/- 100 kb are shown.

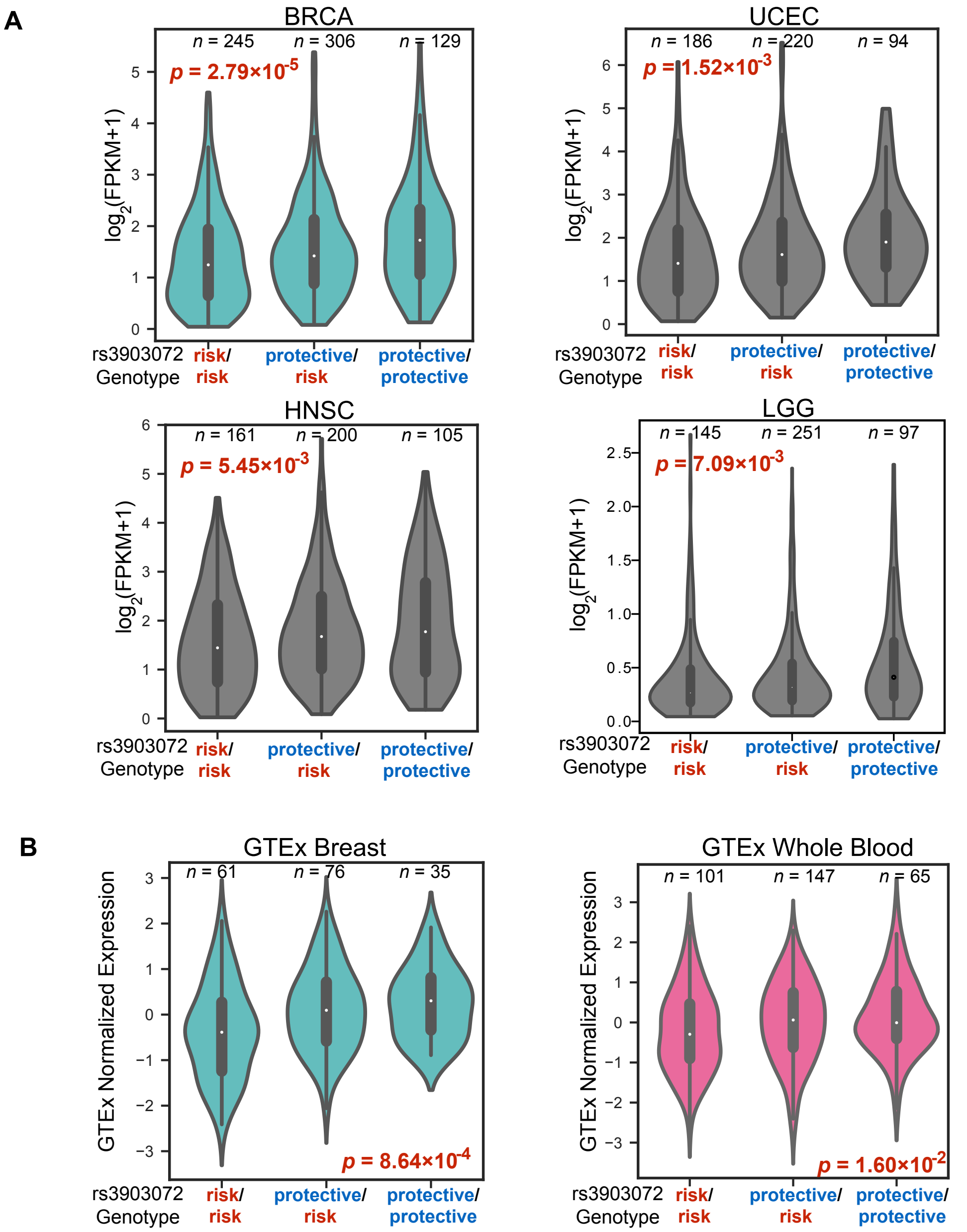

**Supplementary Figure 2.** Violin plots for the eQTL analysis in different datasets. A linear model was constructed between the *CTSW* expression level and the genotype status at the GWAS SNP rs3903072; the *p*-values shown are for the linear coefficient of genotype. Gene copy number is not included in the model for this figure. (**A**) eQTL analysis in cancers from TCGA, using ER+ breast cancer subtype in BRCA, endometrial cancer (UCEC), head and neck cancer (HNSC), and low grade glioma (LGG). (**B**) eQTL analysis in normal tissues from GTEx, using mammary tissue and whole blood tissue.

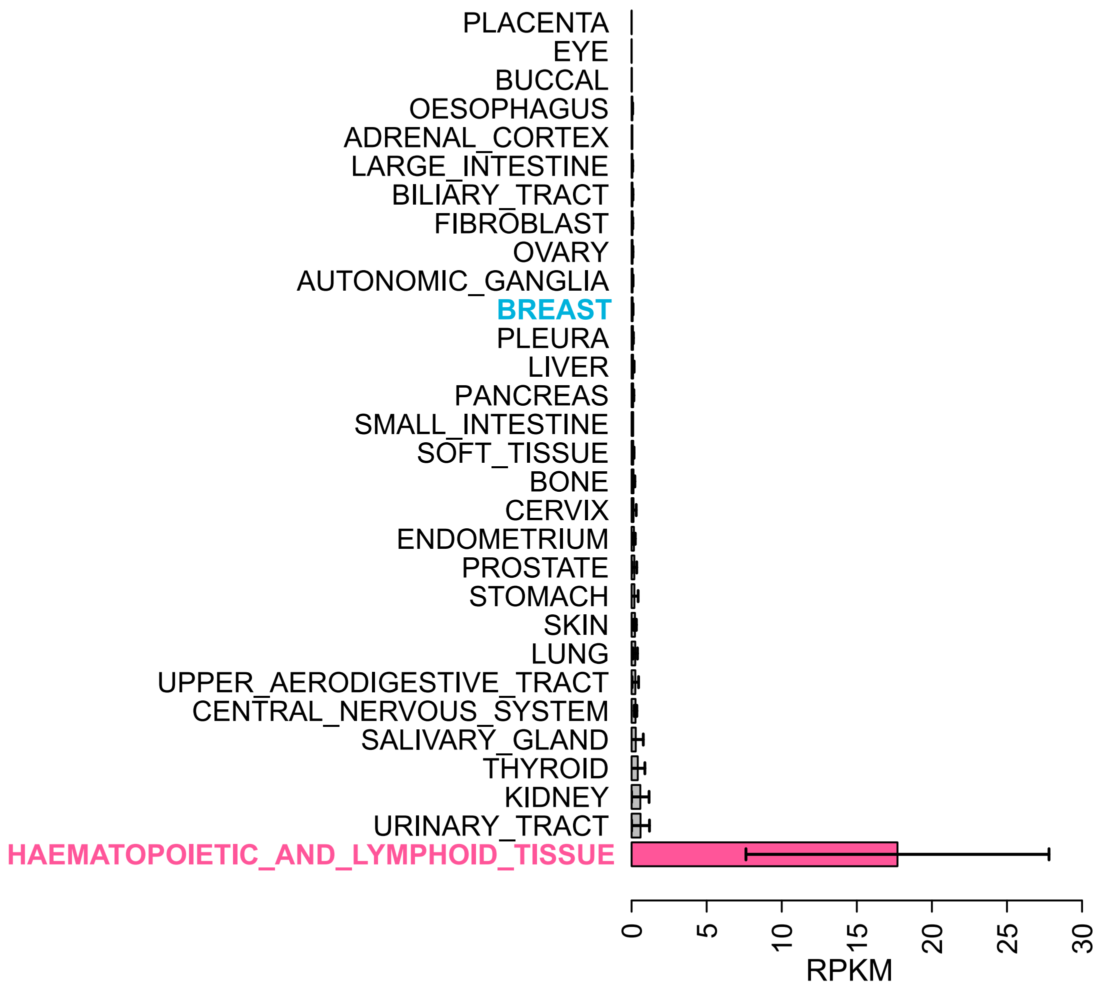

**Supplementary Figure 3.** *CTSW* expression across different cell lines using data from CCLE. Cell lines are grouped by their tissue type; tissues are ranked based on the mean *CTSW* expression of cell lines within each group. An error bar is also shown for each group indicating standard deviation.

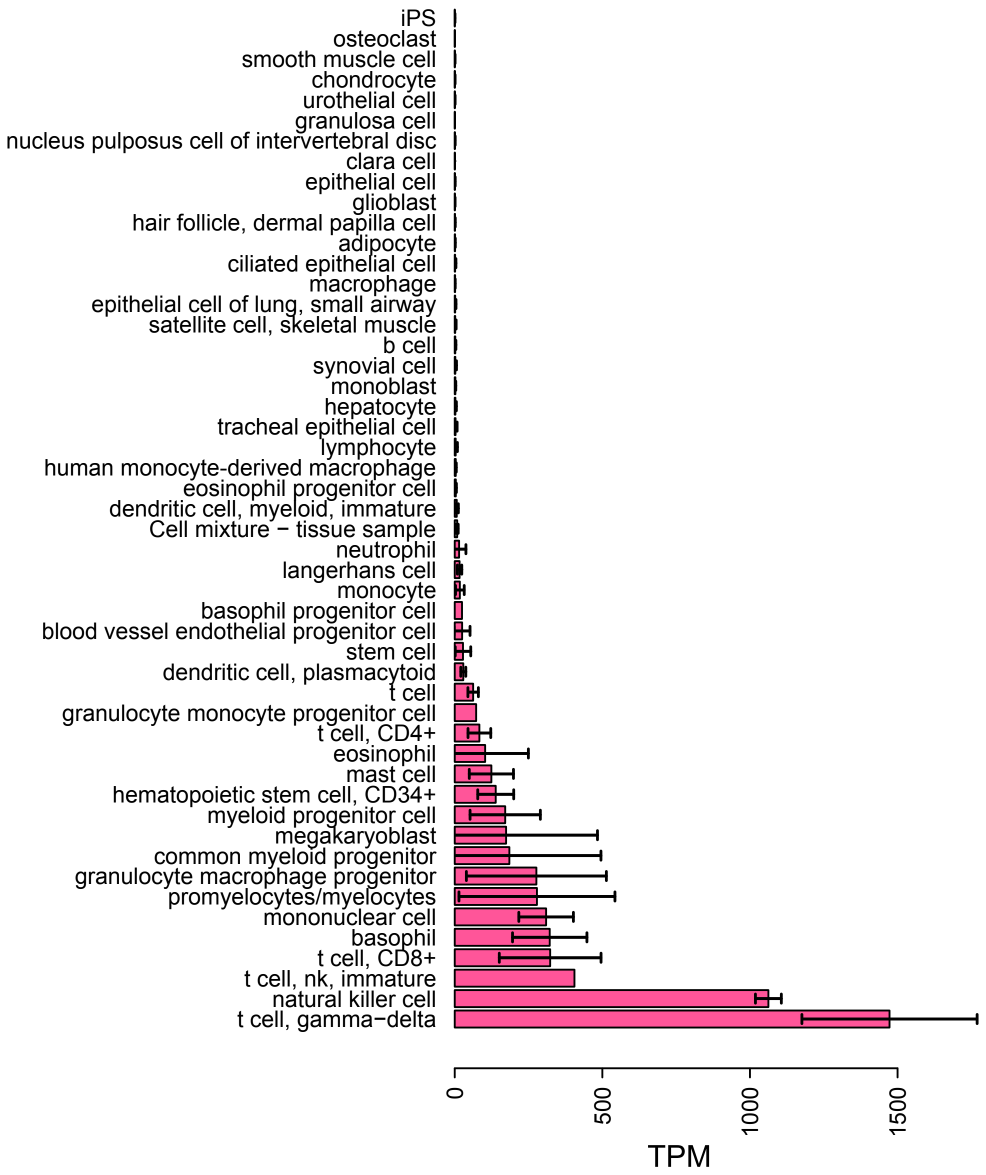

**Supplementary Figure 4.** The *CTSW* promoter transcription activity measured by FANTOM5 for different cell types. Top 50 cell types are shown, ranked by mean *CTSW* expression within each cell type. The cell types listed are obtained from an annotation file provided by the FANTOM5 consortium. An error bar is also shown for each cell type indicating standard deviation. TPM, tags per million.

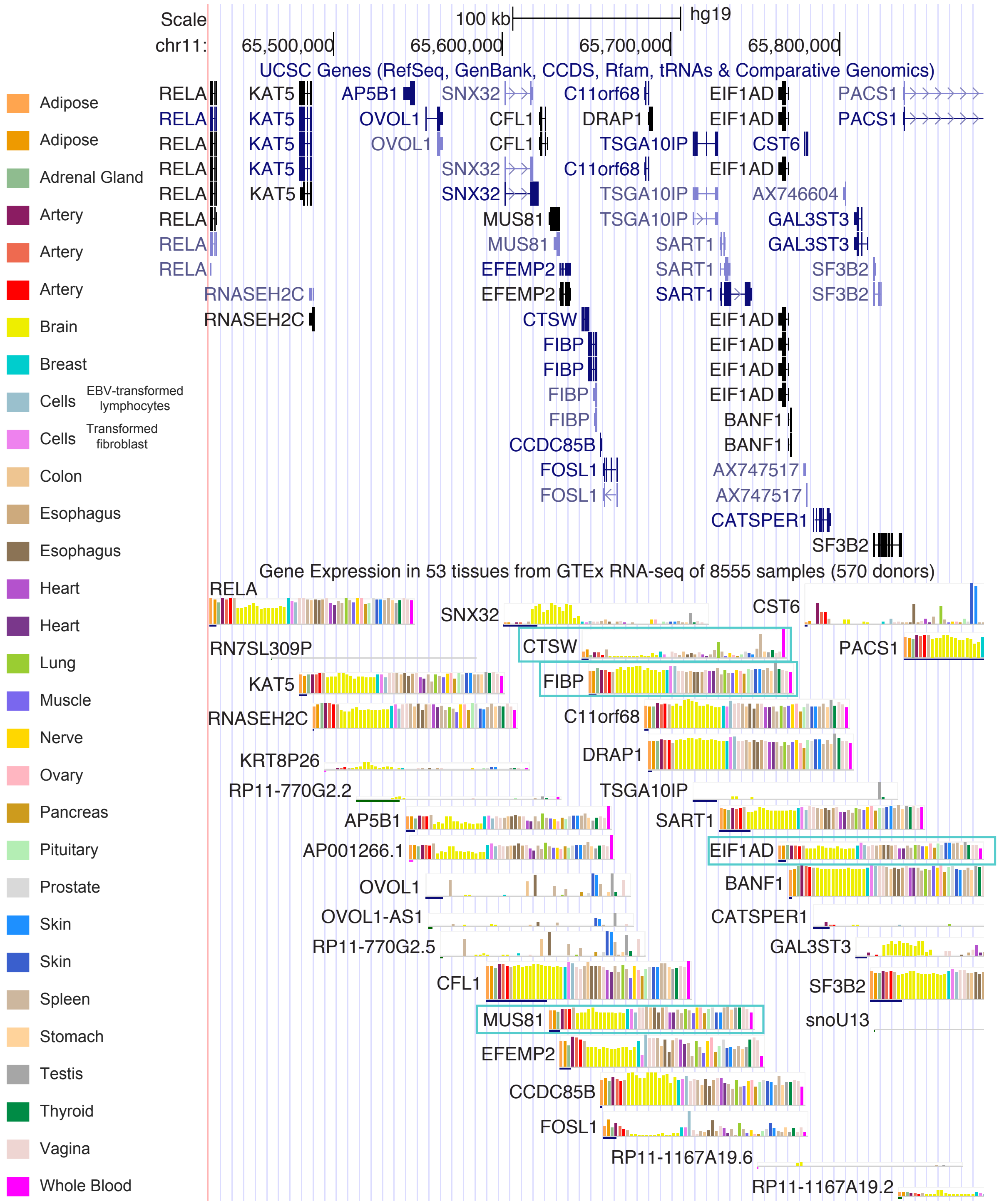

**Supplementary Figure 5.** Tissue-wise expression of *CTSW* and nearby genes. The four eQTL genes shown in Figure 1B (*MUS81, CTSW, FIBP, EIF1AD*) are boxed. The tissue specificity of *CTSW* expression is clearly visible.

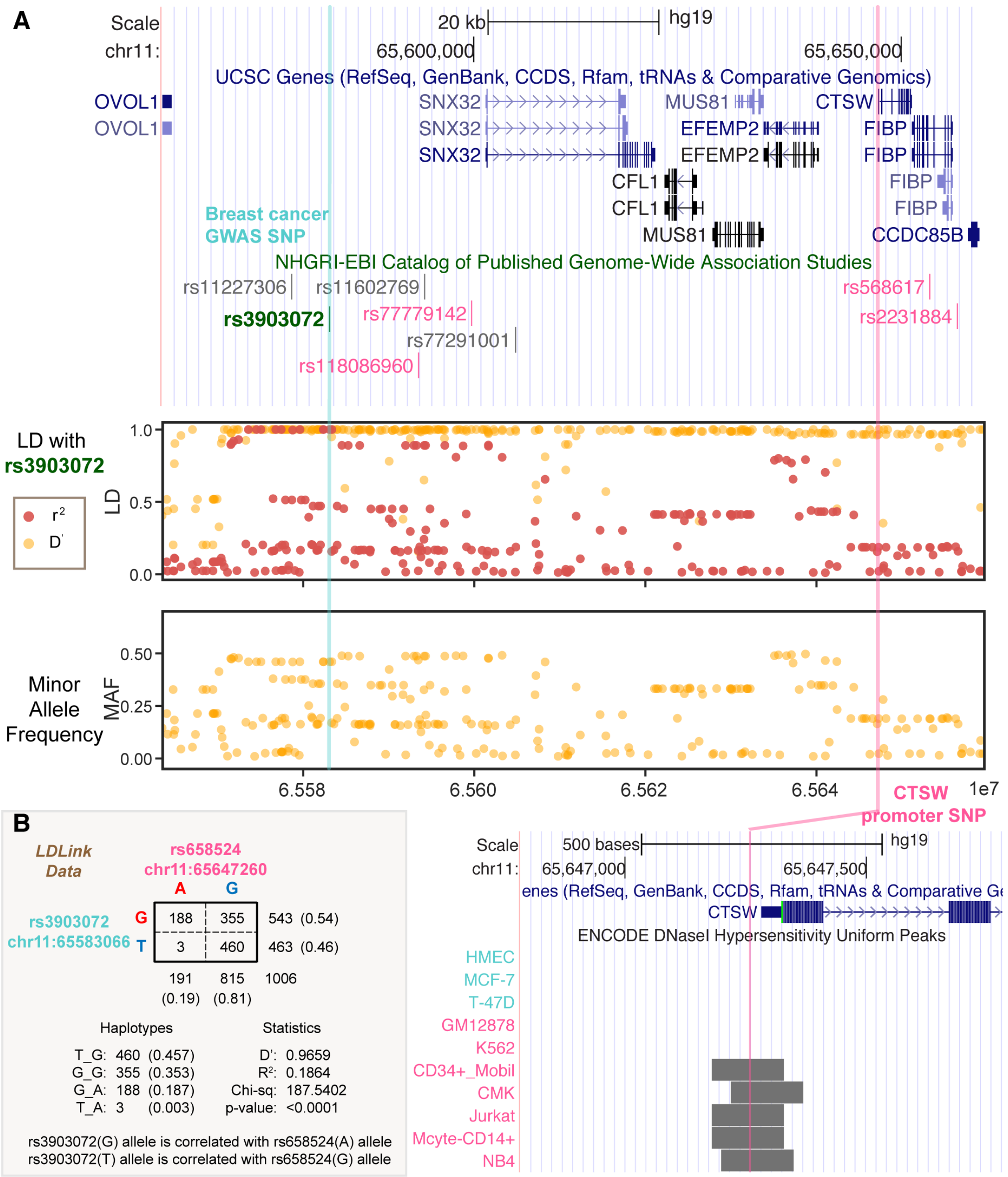

**Supplementary Figure 6.** The putative regulatory SNP in *CTSW* promoter in weak linkage with the GWAS SNP. (**A**) The *CTSW* promoter SNP is in high $D'$ but low $r^2$ with the breast cancer GWAS SNP rs3903072, and it is a rarer SNP compared to rs3903072. Enlarging the *CTSW* promoter region shows that the promoter SNP is located near the center of the *CTSW* DHS in several blood cell lines. This is a zoomed-in region of **Figure 1A** where the immuno-inflammatory GWAS variants are marked magenta, with one more Rosacea SNP rs77779142 (**Supplementary Table 1**). (**B**) The linkage structure between the *CTSW* promoter SNP and the GWAS SNP. Most haplotypes carrying the rs658524-A allele have the rs3903072-G (risk) allele (data presentation from LDLink: https://ldlink.nci.nih.gov, computed based on 1000 Genomes phase 3 EUR population).

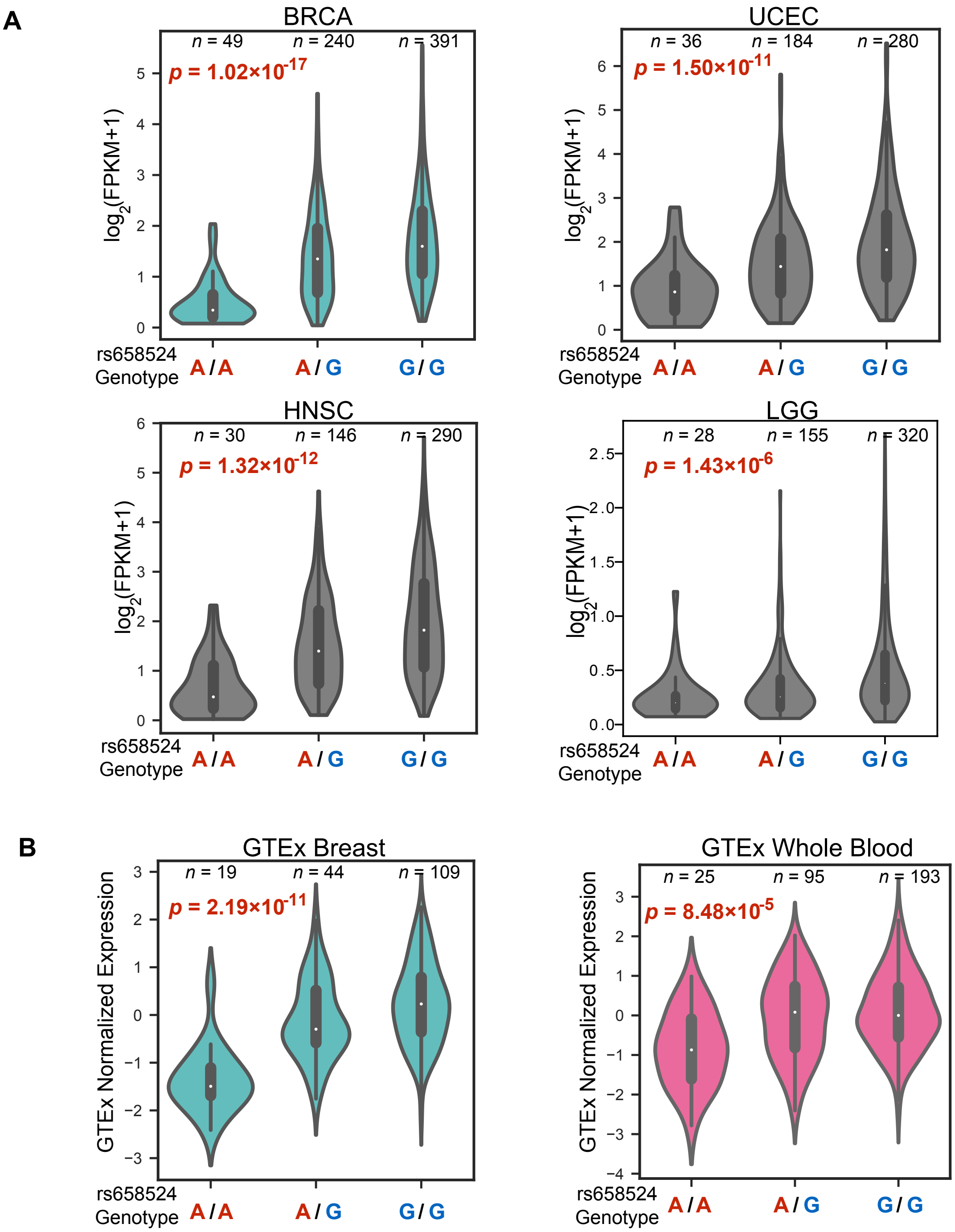

**Supplementary Figure 7.** Violin plots for the eQTL analysis in different datasets, similar to **Supplementary Figure 2** but using the genotypes at the *CTSW* promoter SNP. A linear model was constructed between the *CTSW* expression level and the genotype status at the promoter SNP rs658524; the *p*-values are for the linear coefficient of genotype, and gene copy number is not included in the model for this figure. Note that rs658524-A is marked as being the risk allele (red), as inferred through its haplotype structure with rs3903072. (**A**) eQTL analysis in cancers from TCGA, using ER+ breast cancer subtype in BRCA, endometrial cancer (UCEC), head and neck cancer (HNSC), and low grade glioma (LGG). (**B**) eQTL analysis in normal tissues from GTEx, using mammary tissue and whole blood tissue.

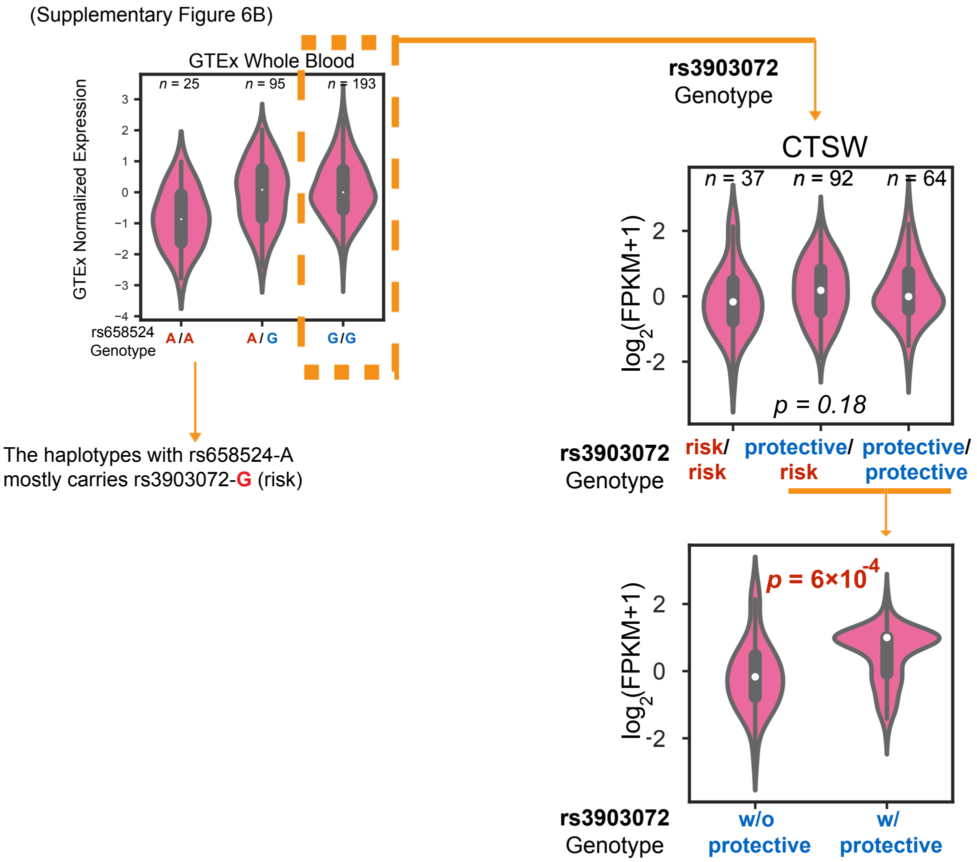

**Supplementary Figure 8.** eQTL analysis conditioned on the protective homozygous genotype at the *CTSW* promoter SNP. We selected the samples carrying G/G at the promoter SNP, and studied whether there exists a residual effect from the distal GWAS genotype on the *CTSW* expression level. A recessive effect of the GWAS SNP is observed in the bottom plot. The *p*-value in the top right plot is for the genotype coefficient from the eQTL linear model, and the *p*-value in the bottom plot is computed using two-sided Welch's t test.

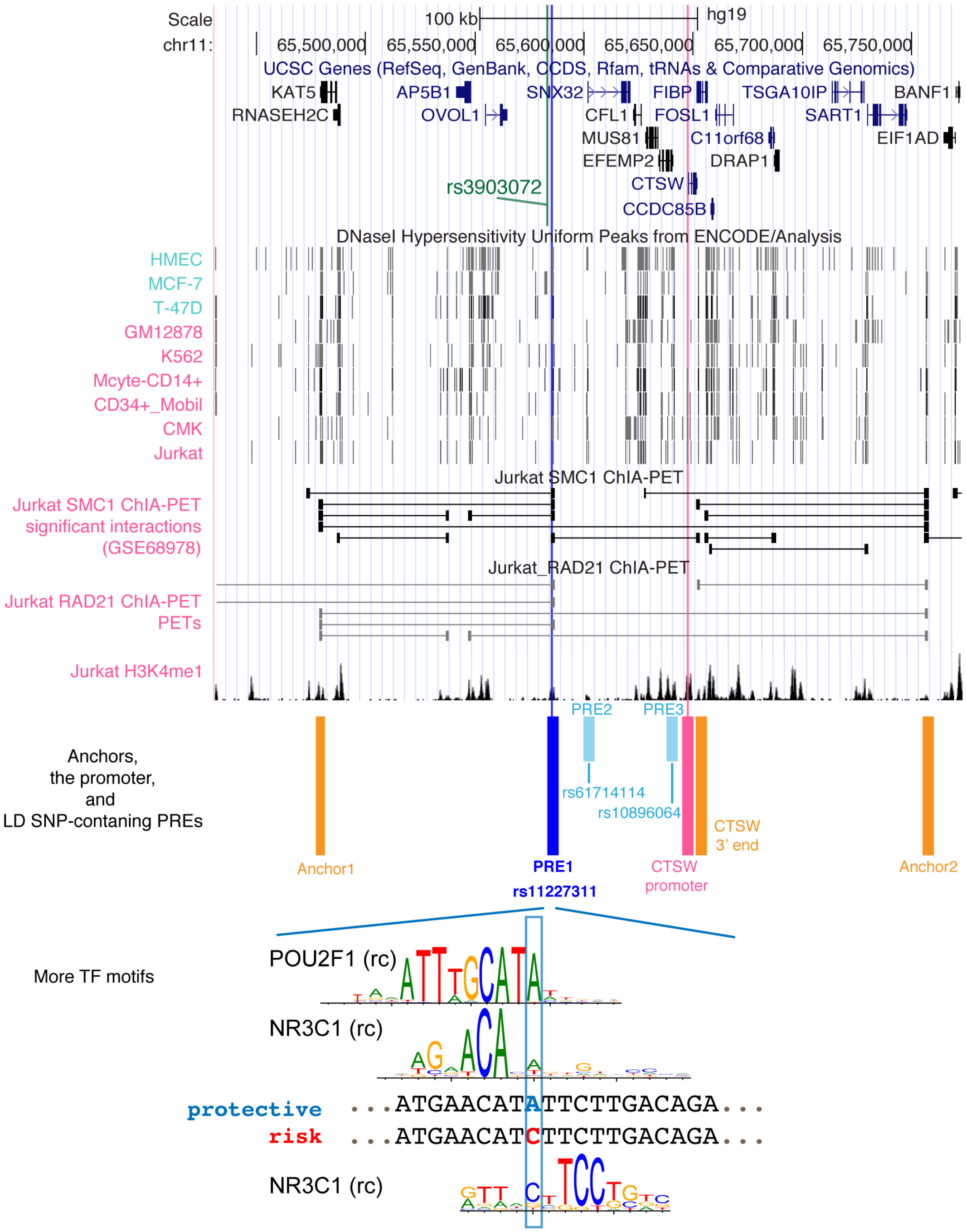

**Supplementary Figure 9.** ChIA-PET data and anchoring elements around the GWAS region. The same region as in **Figure 1A** is shown, presenting DHS tracks, ChIA-PET data in the Jurkat cell line, putative regulatory elements, and anchoring elements that may mediate multi-way chromatin interactions. The putative regulatory elements (PRE), PRE1, PRE2, and PRE3, all contain GWAS-linked SNPs overlapping a blood cell DHS (**Supplementary Table 5**), but PRE1 is highlighted, because of its interaction with the 3' end of *CTSW*. Several possible TF motifs affected by the PRE1 SNP, besides those displayed in **Figure 3B**, are shown here.

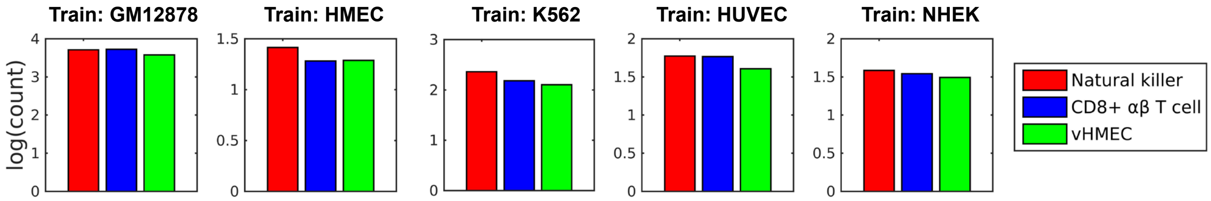

**Supplementary Figure 10.** Predicted log contact counts for the pair between rs3903072-PRE1 and *CTSW* promoter in three cell lines: Natural Killer cells, CD8+ αβ T cells and vHMEC.
